# Brain-like border ownership signals support prediction of natural videos

**DOI:** 10.1101/2024.08.11.607040

**Authors:** Zeyuan Ye, Ralf Wessel, Tom P. Franken

**Author notes:** Correspondence and requests for materials should be addressed to T.P.F.

## Abstract

To make sense of visual scenes, the brain must segment foreground from background. This is thought to be facilitated by neurons in the primate visual system that encode border ownership (BOS), i.e. whether a local border is part of an object on one or the other side of the border. It is unclear how these signals emerge in neural networks without a teaching signal of what is foreground and background. In this study, we investigated whether BOS signals exist in PredNet, a self-supervised artificial neural network trained to predict the next image frame of natural video sequences. We found that a significant number of units in PredNet are selective for BOS. Moreover these units share several other properties with the BOS neurons in the brain, including robustness to scene variations that constitute common object transformations in natural videos, and hysteresis of BOS signals. Finally, we performed ablation experiments and found that BOS units contribute more to prediction than non-BOS units for videos with moving objects. Our findings indicate that BOS units are especially useful to predict future input in natural videos, even when networks are not required to segment foreground from background. This suggests that BOS neurons in the brain might be the result of evolutionary or developmental pressure to predict future input in natural, complex dynamic visual environments.

## Introduction

To understand the world around us, we parse incoming visual information into an organized collection of objects. In primate animals, this capability is thought to be facilitated by neurons in the early areas in visual cortex that encode border ownership (BOS)^1–4^. These neurons fire more to an identical border in their classical receptive field (cRF) depending on which side owns the border, even though the contextual information that defines the side of foreground occurs far outside of the cRF (Figure 1A). This selectivity extends to natural images^5,6^ and the preferred side of ownership corresponds to the side that is near when varying depth^7^. Psychophysics and imaging studies support that BOS neurons also exist in the human brain^8–11^. It is unknown under which conditions BOS signals emerge in neural networks. Artificial neural networks (ANNs) are a great tool to study such ‘why’ questions of how the brain works, because they enable to test whether a particular neural phenomenon results from optimization for a specific task^12^.

**Figure 1.**
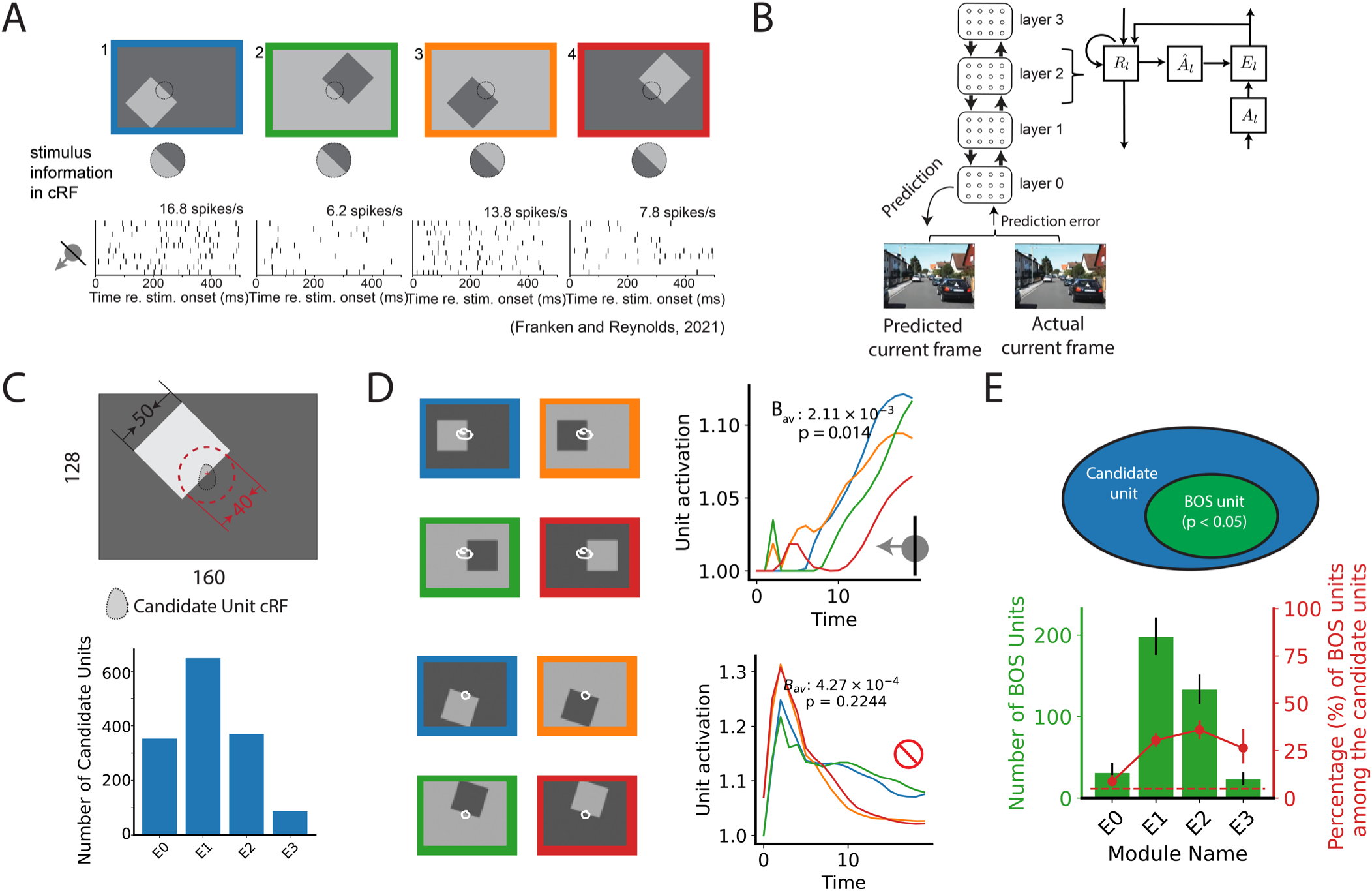
Border ownership (BOS) signals emerge in PredNet. **(A)** An example unit in the primate visual cortex that is selective for BOS. The unit has different responses depending on the BOS, even though the image pixels in its classical receptive field (cRF) are identical for panel 1 and 2, and for panel 3 and 4. The preferred side of ownership is the same for borders in the cRF with a different contrast polarity (the unit fires more to scene 1 than to scene 2, and more to scene 3 than to scene 4). Arrow on the bottom left indicates the side of BOS that this unit prefers. Figure adapted from Franken and Reynolds^3^. (**B**) PredNet is an artificial neural network designed for video prediction. At each time step, the model operates by updating unit activities sequentially from the top layer (layer 3) to the bottom layer (layer 0), generating a prediction of the current video frame. The prediction error is then fed forward to layer 3. Each layer contains four modules (Â_*l*_, *A*_*l*_, *E*_*l*_, and *R* where *l* = 0, 1, 2, 3 indicates layer index, see Methods) (**C**) Candidate units in PredNet are defined as units whose cRF overlaps with the central border but not with any of the square’s corners (see Methods). Bottom: the number of candidate units in *E* modules across different layers. See SI Figure 2 for *R* module data. (**D**) Responses of two example units (module *E2*), with white contours indicating the cRF. *B*_*av*_ measures, for each unit, the selectivity for BOS across different square orientations (see Methods). Colored lines indicate the response to the different stimulus conditions (colors indicate for each response function to which of the stimulus panels on the left it corresponds) for one orientation. *p* value (two-tailed) was computed by comparing *B*_*av*_ to that obtained by shuffling the BOS labels (permutation test, see Methods). BOS units are defined as those units for which this value is smaller than 0.05. (**E**) Number and percentage of BOS units in E-module in different layers. Error bars indicate 95% confidence interval. Horizontal dashed line indicates the chance level for the percentage of BOS units (5%).

It seems intuitive to hypothesize that BOS signals emerge in ANNs when they are explicitly trained on scene segmentation, given that this is assumed to be the primary role of such signals in the brain^13^. A recent study indeed found that units selective for BOS occur in a supervised ANN trained to segment handwritten digits (a processed MNIST dataset^14^) from background^15^. However, such supervised learning has been criticized as biologically highly implausible because it requires a large number of explicitly segmented labels which is unrealistic in brain development^16,17^. Another study found that BOS signals can arise in an unsupervised ANN trained to develop translation invariance for an object presented in isolation, but this mechanism failed for scenes with more than one object^18^, as opposed to BOS signals in the brain^19^. Furthermore, these ANNs can only process simple artificial datasets, unlike neural networks in modern deep-learning frameworks or the human brain, which are high performing on realistic natural visual inputs^20–22^. It thus remains poorly understood when BOS signals emerge in neural networks.

Certain properties of BOS neurons in the brain suggest that BOS signals may be important under dynamic conditions. BOS signals are known to persist for hundreds of milliseconds when the contextual information that defines the side of ownership disappears, as long as the border in the cRF, which has then become ambiguous for BOS, remains^23,24^. Furthermore, these persistent BOS signals can be transferred to other neurons if the ambiguous border lands in their cRF after an eye movement^25^. This hysteresis may make it easier to make sense of dynamic visual input by providing spatiotemporal contiguity.

These observations motivated us to study whether BOS signals emerge in an artificial neural network trained to predict future visual input for natural videos. We studied PredNet, a deep neural network with an architecture inspired by predictive coding^17,26,27^. PredNet was trained on a dataset of natural videos captured by car-mounted cameras (KITTI^28^) to predict the next video frame. Our *in-silico* experiments demonstrate that a significant fraction of units in PredNet exhibit BOS signals. Moreover, these BOS units share several properties with BOS neurons in the brain. Finally, ablating PredNet’s BOS units increased prediction error more than ablating the same number of non-BOS units. BOS units thus contribute to prediction of natural visual input even if there is no need to segment foreground from background. This suggests that the need to predict future input in natural videos may drive the development of BOS neurons. These scene segmentation signals, typically considered an example of a ventral ‘what’ stream operation, may thus be more involved in processing dynamic aspects of visual input than is typically assumed.

## Results

### BOS signals emerge in an artificial network trained to predict the next frame of natural videos

To study the role of BOS signals in the processing of complex dynamic input we employed PredNet, a hierarchical ANN introduced by Lotter et al.^26^ (Figure 1B). PredNet comprises four layers, with four modules per layer: the representation (*R*_*l*_), the predicted output (Â_*l*_), the prediction target (*A*_*l*_), and the prediction error (*E*_*l*_) modules. At each time step (see Methods), signals propagate from the top layer to the zeroth layer, resulting in a prediction for the next video frame in Â_0_. This prediction is then compared to the actual next frame provided in *A*_0_. The prediction error signal subsequently propagates from the zeroth layer to the top layer. The network was trained to minimize the prediction error of videos in the KITTI dataset^28^, which were captured by car-mounted cameras in various urban and rural settings in Germany.

We tested whether the BOS units exist in the PredNet by doing an *in-silico* experiment that is analogous to the neurophysiological studies on BOS^1,3,29^ (Figure 1A). We measured the cRF of each unit using sparse noise stimuli (SI Figure 1). First we identified candidate units for BOS tuning. For a unit to be a candidate unit, the cRF needed to include the center of the square’s central border (i.e. the border positioned at the scene center) but must exclude any other border of the square (Fig. 1C; Methods). This criterion is similar to that used in neurophysiological studies on BOS^1,3^. We found tens to hundreds of candidate units in different PredNet modules (Fig. 1C and SI Figure 2). We then analyzed the response from candidate units to the standard full square stimuli. Figure 1D shows responses from two example candidate units. The top unit exhibited a larger response when the square was positioned on the left side of the central vertical border compared to the right side, irrespective of the contrast polarity across the border (i.e. blue vs. green, and orange vs. red). This unit thus prefers that the border in its cRF is owned by a square on one side, similar to BOS neurons in the primate visual cortex^1,30^. In contrast, the bottom unit did not exhibit a clear difference: its response was very similar for stimuli with opposite border ownership but identical contrast polarity of the central border. To quantify BOS tuning for each unit, we first computed the unit’s difference in response between stimuli of opposite border ownership across different contrast polarities, and divided it by the sum of the responses, resulting in the BOS index (BOI). The BOS index was then averaged across all square orientations, resulting in *B*_*av*_ (see Methods). The statistical significance of *B*_*av*_ was determined by comparing it to a null distribution obtained by shuffling the stimulus labels. A candidate unit with a p-value smaller than 0.05 was defined as a BOS unit; otherwise, it was defined as a non-BOS unit. The top unit in Figure 1D has a statistically significant *B*_*av*_ and is therefore a BOS unit, while the bottom unit is a non-BOS unit.

**Figure 2.**
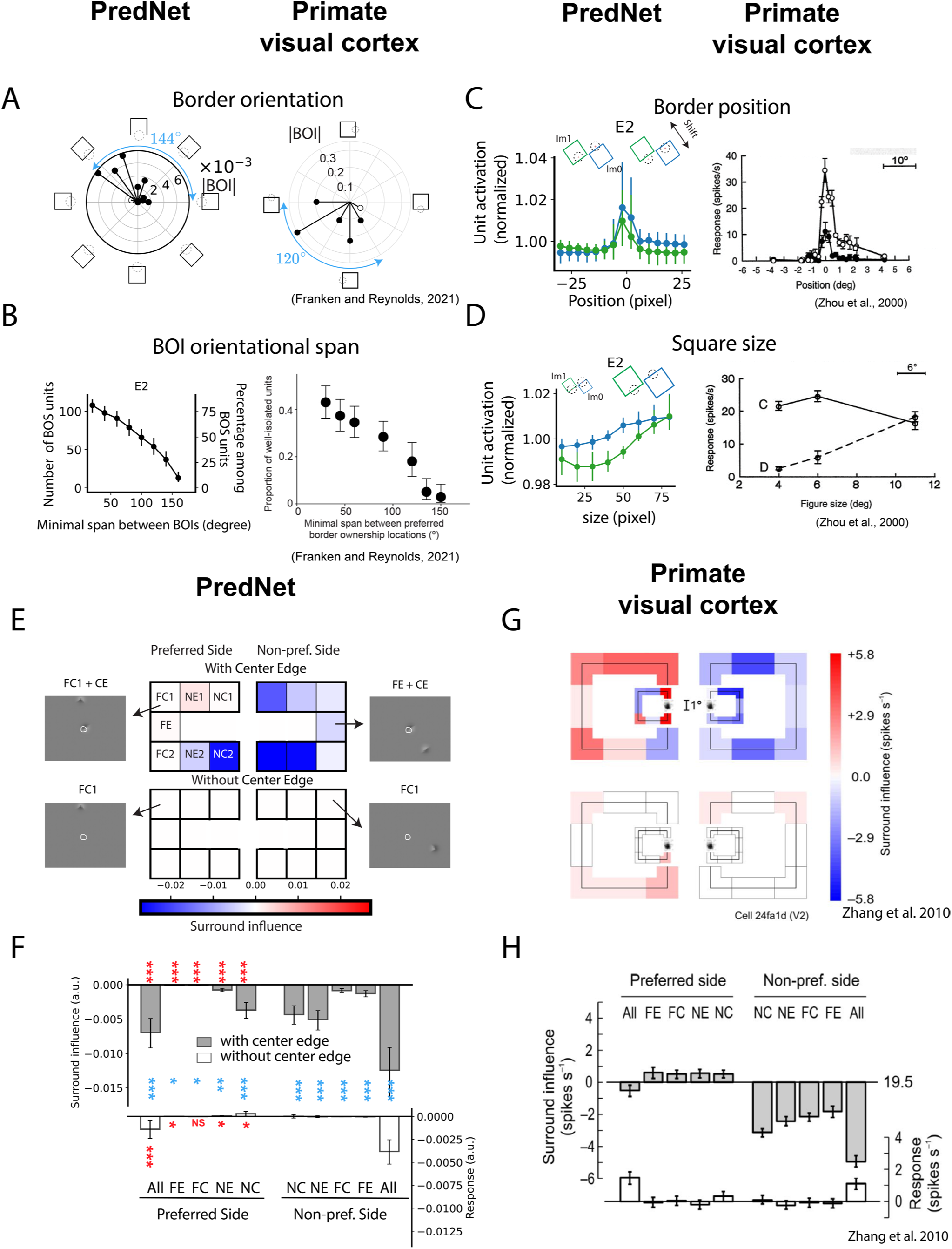
BOS units in PredNet share several properties with BOS neurons in the brain. **(A)** Border ownership index (BOI) at different orientations. Vector magnitude represents the absolute value of BOI, i.e. the difference in unit response to scenes with squares that share a border with a given orientation, but for which the square is positioned on opposite sides of that border (thus a pair of stimulus cartoons on opposite sides of the polar plot), divided by their sum (see Methods). Vector angle is such that the vector points towards the stimulus cartoon with the preferred side of BOS for that border orientation. Left: example of a BOS unit in PredNet. Right: BOS neuron recorded in macaque area V4 (reproduced from Franken and Reynolds 2021^3^). Filled symbols in both panels indicate for which orientations the BOI was significantly different from 0. Blue text indicates the span of a unit, which is the angle between the preferred object locations for orientations with statistically significant BOI (Methods). **(B)** The y-axis illustrates the number/percentage of BOS units whose spans equal or exceed the span indicated by the x values. Error bars indicate 95% confidence intervals. Left: BOS units in module *E*_2_in PredNet (see SI Fig. 4A for other modules). Right: population data from BOS neurons in macaque area V4 (reproduced from Franken and Reynolds 2021^3^). **(C)** Left: for each BOS unit, squares were generated with different positions as indicated in the cartoon. The blue and green traces represent normalized population responses (see Methods) to opposite BOS (blue corresponds to the preferred BOS derived from responses to the standard square set). Dots and error bars show the median, first, and third quantiles across the population of BOS units. Right: responses from a BOS neuron in macaque V2 for different square positions. The two traces indicate opposite BOS. Dots and error bars represent mean firing rates and SEMs across trials (reproduced with permission from Zhou et al. 2000^1^. Copyright 2000 Society for Neuroscience). **(D)** Identical to (C) but square size was varied instead of square position. Right panel reproduced with permission from Zhou et al. 2000^1^. Copyright 2000 Society for Neuroscience. (**E**) Response of an example BOS unit in PredNet (module *E*_2_) to square fragments. Top half shows responses to a square fragment in the surround paired with the border in the cRF. Bottom half shows responses to square fragments in the surround without the border in the cRF. Gray panels show example scenes (white outline: cRF). Colors of the central panels indicate the surround influence. The surround influence is the unit’s response to a scene with a square fragment in the surround at the position indicated by the letter codes (also symbolized by the panel’s position), subtracted by that to a scene without the square fragment. Letter codes: NC: near corner; NE: near edge; FC: far corner; FE: far edge; numbers indicate different positions of the fragment, e.g. NC1 and NC2 refer to each of the near corners on opposite ends of the central border. (**F**) Means and 95% confidence intervals (i.e. 1.96 times SEM) of surround influence across all BOS units in module *E*_2_ (n=71; ; see SI Fig. 5 for other modules). NC is the average of NC1 and NC2, and the same was done for NE and FC. ‘All’ represents the surround influence when all square segments were shown (top), or all square segments except the center edge were show (bottom). Red text indicates whether the surround influence for a particular condition is significantly larger on the preferred side than on the non-preferred side. Blue text indicates whether the absolute value of surround influence of with-CE scenes is significantly larger than without-CE scenes. Wilcoxon signed-rank test. ***: p < 0.001; **: p < 0.01; *: p < 0.05; NS: no significance. Outlier units (see Methods) were removed to compute mean and SEM but included in the statistical tests. (**G**) Same as (E) for a BOS neuron in the macaque visual cortex (reproduced with permission from Zhang et al. 2010^31^). Two different square sizes were evaluated, for which surround influence is plotted separately as the smaller and larger panels. (**H**) Similar panel as (F), for BOS neurons in the macaque visual cortex, with permission from Zhang et al. 2010^31^.

We conducted a population analysis of *B*_*av*_ across all candidate units. We find that 20-40% of candidate units in *E_1_*, *E_2_* and *E_3_* have significant *B*_*av*_values (Fig. 1E). This is larger than in module *E*_0_ (8.7 %, 95% confidence interval [6.2%, 12%]). BOS units were also found in modules *R_1_* and *R_2_*, but much less in *R_0_* (*R_3_* had only a small number of candidate units; SI Fig. 2). The distribution of BOS units in PredNet’s hierarchy is reminiscent of the distribution of BOS neurons in the primate visual cortex, which are less prevalent in areas closer to the sensory input (V1) than in downstream areas (V2 and V4)^1,3,4^.

### PredNet’s BOS signals are robust to scene variations common in natural object transformations, like BOS neurons in the brain

We explored the robustness of BOS signals to the same scene variations that have been used in neurophysiology studies on BOS neurons: square orientation, position, and size^1,3^. Figure 2A (left) shows the BOI for different square orientations for an example BOS unit. Vector length indicates the absolute value of the BOI, and the angle of each vector indicates the preferred side of BOS for each orientation (cf. the symbols around the plot). The square orientations with a large BOI form a contiguous region in visual space, which is similar to BOS neurons in the primate visual cortex (e.g. Fig. 2A, right)^3^. Filled symbols in Fig. 2A indicate orientations for which BOI is significant (permutation test, see Methods). The angular span between object locations at the preferred side of BOS for different border orientations (only orientations for which BOIs is significant are considered) is referred to as the BOS span. For example, the span for the neuron shown in Figure 2A (left panel) is 144°. A substantial number of BOS units in PredNet have a large span, extending to ∼150°, similar to BOS neurons in the brain (Fig. 2B; SI Fig. 4A).

Next, we examined BOS tuning for different square positions and sizes. We set the orientation at that for which |BOI| was maximal, and then varied position (the position of the center of the central border varied along a line orthogonal to the border’s orientation; Fig. 2C). We also varied square size for the central position (Fig. 2D). We find that the response difference between scenes with opposite BOS was consistent for different positions or sizes in the population of BOS units in PredNet (Fig. 2C,D, left panels), just like for BOS neurons in the brain (Fig. 2C,D, right panels)^1^. We quantified this consistency by averaging BOI across different conditions (i.e., size or position, see Methods). For all modules with a substantial number of BOS units (over 15 units), these averaged BOI values are statistically significantly positive (i.e. consistent with the tuning in the baseline condition; SI Figure 4B, C, bootstrapping test, see Methods). Taken together, we find that border ownership signals in PredNet are robust to differences in square orientation, size and position, i.e. remarkably similar to BOS neurons in the primate visual cortex^1,3^.

### Surround influence for PredNet’s BOS units is similar to BOS neurons in the brain

Neurophysiological experiments found that isolated object fragments in the surround modulate the activity of BOS neurons in a way that is consistent with BOS tuning: fragments on the non-preferred side of BOS suppress the response significantly more than fragments on the preferred side, which often have an enhancing effect^31^. These modulatory effects were only significant in the presence of a border in the cRF. We analyzed how fragments in the surround modulated the activity of BOS units in PredNet. Similar to Zhang et al. 2010^31^, we divided a square object into 8 fragments: one Center Edge (CE) located at the image center, and 7 contextual fragments (two Near Corners [NC1 and NC2], two Near Edges [NE], two Far Corners [FC] and one Far Edge [FE]). This allows us to create two types of fragment scenes. The first type pairs one of the fragments in the surround with the CE (‘with-CE’). Two additional scenes contain respectively only the CE, or all the fragments. The second type are identical scenes but without the CE (‘without-CE’). For ‘with-CE’ scenes we defined the surround influence of a fragment as the unit’s response to the combination of that fragment and the CE, subtracted by the response to the CE-only scene (see Methods). For ‘without-CE’ scenes, the surround influence of a fragment was defined as the response to a scene with that fragment, subtracted by the response to a full-gray scene. Figure 2E displays the data for one example BOS unit in PredNet. First, we noticed that the absolute value of surround influence in ‘with-CE’ scenes is larger than in ‘without-CE’ scenes. This was the case for each PredNet module with at least 10 BOS units (Fig. 2F for *E*_2_ and other modules in SI Fig. 5). This is similar to BOS neurons in the visual cortex (Figs. 2G,H)^31^. Second, we compared the modulation effect between fragments on the preferred side and the non-preferred side. The preferred and non-preferred sides were determined solely from the responses to standard square scenes (Fig. 1A). Despite this, we found that for all modules with more than 10 BOS units, the surround influence for most fragments is significantly more negative when they are presented on the non-preferred side compared to the preferred side (Fig. 2F; SI Fig. 5). This is similar to BOS neurons in the visual cortex (Figs. 2G, H). These data indicate that BOS tuning in PredNet does not result from a single hotspot in the surround, but that multiple fragments collectively contribute, as is the case for BOS neurons in the brain^31^.

### PredNet’s BOS units exhibit hysteresis, similar to BOS neurons in the brain

A remarkable characteristic of BOS neurons in the brain is that the BOS signal persists for hundreds of milliseconds, even when the contextual information that defines the side of ownership disappears^24,25^. We tested if BOS units in PredNet also exhibit this phenomenon. We used a Square-Ambiguous sequence similar to what was used in physiology experiments^24^. The sequence consists of a full square scene (Figure 1A) in the first four time steps, which transitions into a scene with a border that is ambiguous for border ownership (Figure 3A, left). We presented these sequences to PredNet and computed the time course of the relative response difference (RRD), defined as the difference in response between the preferred and non-preferred square sides, normalized by the average response (see Methods). The RRD for BOS units remains positive for multiple time steps (Fig. 3B left, red function; SI Figure 6). The units thus respond differently to the ambiguous scene (which is identical in the two sequences), depending on stimulus history, a phenomenon called hysteresis. BOS neurons in macaque visual cortex show a similar hysteresis (compare with Fig. 2A in O’Herron and von der Heydt, 2009^24^).

**Figure 3:**
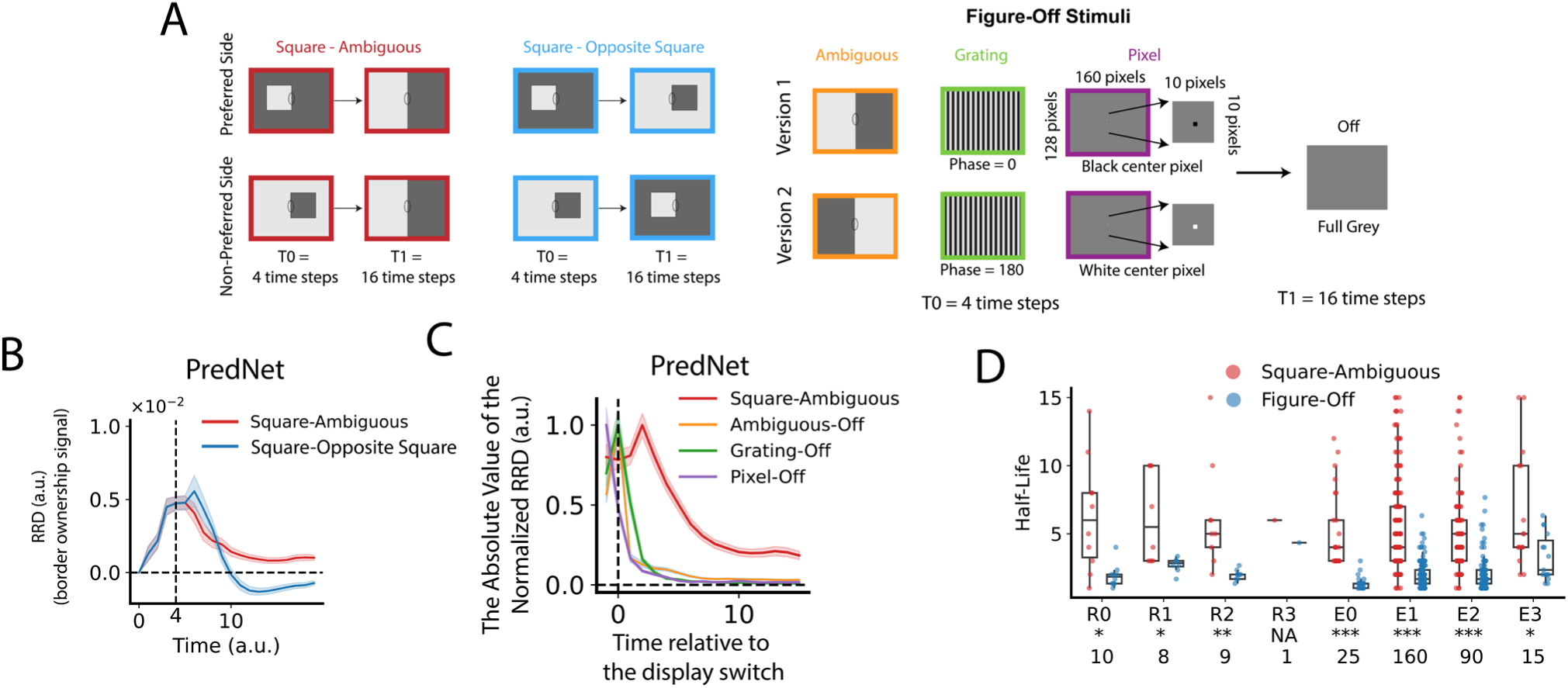
BOS signals in PredNet exhibit hysteresis, similar to BOS neurons in the brain. (**A**) Three sequences with scene changes were used: squares transitioning to ambiguous borders (Square-Ambiguous), squares transitioning to squares with opposite border ownership (Square-Opposite Square), simple scenes (Ambiguous, Grating, or Pixel) transitioning to a full gray scene (Figure-Off). Stimuli were presented at the orientation for which |BOI| was maximal. (**B**) The relative response difference (RRD, see Methods) represents the difference in response between scene sequences that start with a square on the preferred and the non-preferred side (for Square-Ambiguous or Square-Opposite Square), or between version 1 and version 2 (for Figure-Off). Panel shows RRD of BOS units from PredNet (*E*_2_module, n = 132 units). Functions plotted in the same format as Fig. 2A in O’Herron and von der Heydt, 2009^24^. Line and error bands represent the mean and SEM. (**C**) Mean and SEM of the absolute value of the normalized RRD (normalized to maximal value) across BOS units in the *E*_2_ module for different sequences. (**D**) Half-life is defined as the number of time steps after which RRD is reduced to half of its maximum. Each dot corresponds to one BOS unit. Figure-Off data shows the average across the three subtypes shown in A. Only units for which half-life was defined for all conditions were included in this panel (see Methods). Numbers at the bottom indicate the number of included units per module. Asterisks indicate the statistical significance of the difference in half-life between Square-Ambiguous and Figure-Off: NA: not applicable; *p<0.05, **p<0.01, ***p<0.001 (Wilcoxon signed-rank test).

To determine whether this persistent BOS signal is longer than the typical signal decay, we analyzed the response for two control sequences. The first is Square-Opposite Square, which starts with a full square and then switches to another full square image with opposite border ownership (and opposite luminance, so that the contrast polarity of the central border remains the same; Fig. 3A, middle). The RRD for this sequence decays much faster and stabilizes at a negative value, reflecting the switch in BOS (Fig. 3B, left, blue function; SI Figure 6). Again the same pattern occurs in BOS neurons in the brain (compare with Fig. 2A in O’Herron and von der Heydt, 2009^24^). The second control sequence is Figure-Off, in which a simple scene (three subtypes: ambiguous, grating or pixel) is followed by a full gray scene (Fig. 3A, right). Again the RRD of PredNet’s BOS units decays faster to these sequences than to the Square-Ambiguous sequence (Figs. 3C,D, SI Figure 6B). Across all modules with at least 10 BOS units the RRD half-life is significantly longer for Square-Ambiguous sequences than for Figure-Off sequences (Fig. 3D). Together, we find that BOS signals in PredNet have similar dynamic characteristics as BOS neurons in the brain: the BOS signal persists when contextual information disappears such that the side of BOS becomes ambiguous, but quickly updates when the context indicates a switch in BOS.

### BOS units contribute more to prediction than non-BOS units for videos with moving objects

Our data presented thus far demonstrate that units with brain-like tuning for BOS exist in PredNet, a network trained to predict future visual input in video sequences. This suggests that BOS units specifically aid in predicting future video frames. To test that, we conducted ablation experiments in PredNet. We presented Translating-Square videos (40 unique videos in which a square moves at a constant velocity, SI Fig. 7, top) to PredNet. We measured the prediction performance of PredNet to these videos, both before and after ablating either BOS units or non-BOS units (i.e. candidate units that did not pass the criterion for BOS-selectivity, see Methods).

The impact of unit ablation on video prediction is shown in SI Figure 9 (top row). Here, we introduce the metric “relative prediction mean squared error (RPE),” defined as the normalized difference (post-vs. pre-ablation) of the mean squared prediction error (see Methods). A positive RPE represents an increase in prediction error after ablation. To quantify the overall effect of ablation in each module, we measured the slope of the relation between RPE and number of ablated units using linear regression, and a bootstrapping test to assess the statistical significance of this slope between ablating BOS units or non-BOS units (indicated with red symbols in SI Fig. 9, top row). We find that the RPE is significantly higher when BOS units were ablated than when non-BOS units were ablated for most modules. We wondered if this could be explained by a difference in responsiveness: BOS units may respond more to these video frames than non-BOS units. To explore that possibility, we subsampled the populations to ensure there were no statistically significant differences in response magnitude to the videos (Wilcoxon rank-sum test, p > 0.5, see SI Figure 8 and Methods). The ablation experiment in these subsampled populations shows the same pattern, ruling out that the RPE difference is due to a difference in average response (Fig. 4, top row). The data thus indicate that BOS units contribute more than non-BOS units in predicting future frames for these videos.

**Figure 4.**
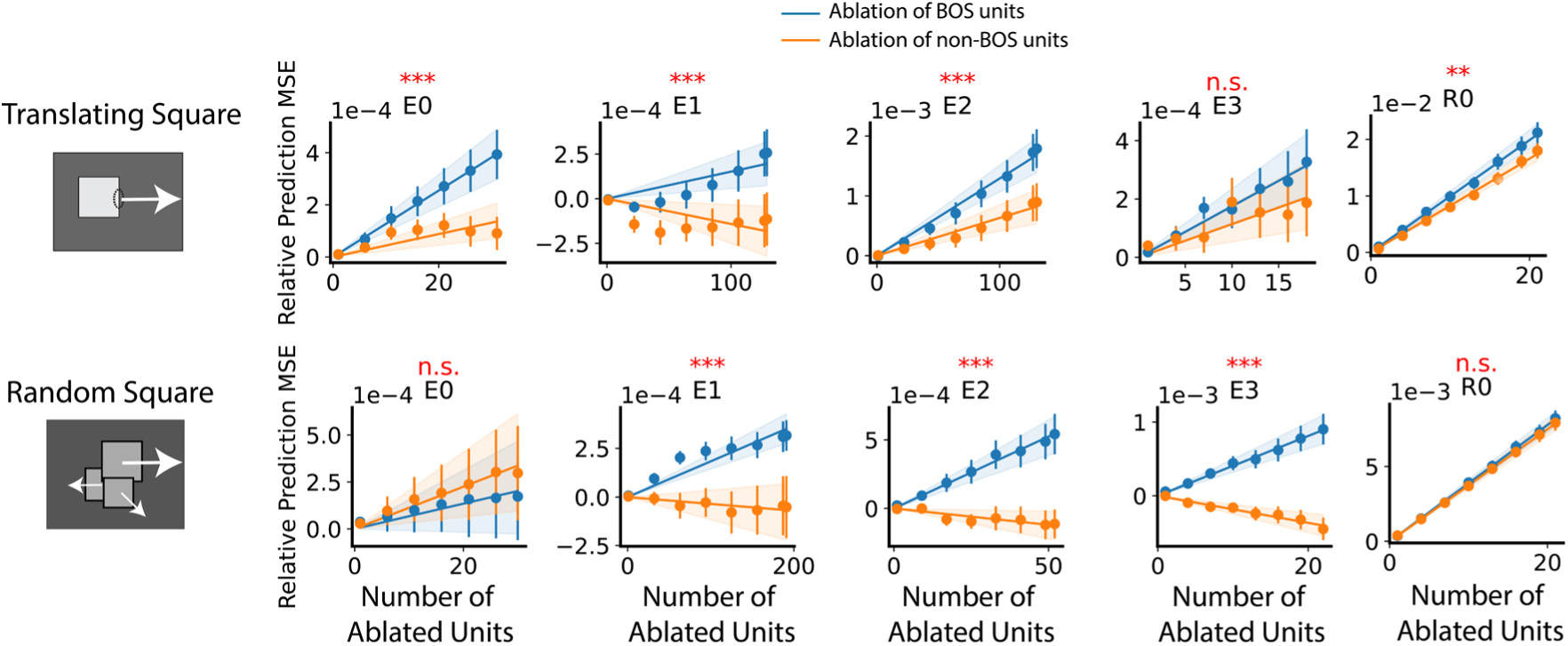
Ablating BOS units in PredNet increases prediction error more than ablating non-BOS units for videos with moving objects. The left shows an example frame of each video type (arrows indicate motion and are not part of the frame). Translating Square videos show a square moving at constant speed; Random Square videos show a random number of squares of different sizes, initialized at random positions and moving at random, constant velocities (see also SI Figure 7). Right panel shows the relative prediction mean squared error (RPE) for different numbers of ablated units. RPE measures the relative change of prediction error due to ablation. Non-BOS units are candidate units that do not pass the criterion of BOS selectivity. Dots and error bars denote respectively the mean and SEM of the RPE across 10 randomly chosen video samples. The RPE of one video sample is the average RPE of 10 samples of unit ablation (Methods). The solid line indicates the best linear fit, with bands indicating the 95% confidence interval. The red text above the panels indicates whether the slopes of the lines differed significantly between BOS- and non-BOS-unit ablation. n.s.: not significant; **: p < 0.01; ***: p < 0.001 (bootstrapping test). Modules *R*_1_, *R*_2_ and *R*_3_ contain a small number of candidate units and are therefore not included in this analysis.

We wondered if BOS units also contribute to prediction of videos with multiple objects. We generated videos with several squares that were randomly positioned, and moved in random directions (SI Fig. 7, middle). When we performed the same ablation experiment for these videos, we find the same pattern: BOS units typically contribute more to prediction than non-BOS units, even though, again, PredNet was not exposed to such videos during training (Fig. 4 bottom, SI Fig. 9 middle).

Finally, we wondered if BOS units aid in prediction with any video. We performed the same experiment in a set of 41 natural videos from the KITTI database^28^ (SI Fig. 7, bottom). This is the same database that was used to train PredNet, but we only included videos that were not used during training. BOS units in the *E*_2_ contribute more to prediction than non-BOS units (SI Fig. 9). Note that these videos are much higher-dimensional than the translating square and random square videos, and there is a high degree of heterogeneity within the small set of 41 videos. This results in smaller overall RPEs when averaged across videos than for the square videos, and not enough statistical power to precisely estimate RPE in the subsets of units with similar responsiveness (SI Fig. 10).

Together these experiments suggest that BOS units emerge in PredNet because they contribute more to prediction than non-BOS units for videos with moving objects.

## Discussion

The assignment of borders to foreground surfaces is thought to be a key step in visual scene segmentation^13,32^, and a substantial fraction of neurons in visual areas V2 and V4 of the primate brain signal this ownership of local borders^1–3^. It is poorly understood why the brain resorts uses this particular representation. Here we discovered that units selective for BOS also emerge in an artificial neural network, PredNet^26^, that was trained to predict future input in natural videos. Importantly, the network was not explicitly trained to distinguish foreground from background or to identify objects in visual scenes. Interestingly, BOS units in PredNet share several properties with BOS neurons in the brain (robustness for different positions, orientations, and sizes^1,3^; asymmetric functional effects of object fragments on opposite sides of the border^31^; BOS hysteresis^24^), suggesting that these signals are functionally similar to those in the brain. Finally, we found that ablation of BOS units affects prediction accuracy more than ablation of non-BOS units. Overall, our results suggest that BOS units might emerge in neural networks trained on natural, complex dynamic input primarily because they are particularly helpful to efficiently process such input, even if segmentation is not required.

PredNet’s architecture was inspired by the predictive coding framework. This theory proposed that a major function of the sensory cortex is to predict incoming sensory stimuli^33–37^. The hierarchical organization of visual cortical areas is proposed to compute an internal model of the external world, and feedback from areas higher in the hierarchy (e.g. V4, IT) is thought to reflect predictions from this internal model, which is then compared with incoming sensory stimuli in lower areas (e.g. V1, V2)^33^. There are hints suggesting that how brain circuits compute border ownership may be understood in this framework^13,38–40^. For example, area V4 has been proposed to contain grouping cells which compute proto-object representations with short latency, i.e. an early prediction of the shape and location of objects in the scene. Feedback from such cells could explain border ownership signals in lower areas^6,13,31,41^, and a recent study indeed found evidence that supports the existence of grouping cells in V4^23^. Response dynamics and laminar organization of BOS neurons align better with feedback models than with alternatives that solely rely on intra-areal horizontal connections or feedforward connections^3,4,31,42–46^. Moreover, the phenomenon of BOS hysteresis indicates that BOS neurons persistently signal the most likely scene organization even if contextual information disappears, but quickly update when sensory information inconsistent with the current internal model appears^24,47^. The present data provide complementary evidence that there may indeed be a tight link between predictive coding and how neural networks compute BOS. We showed that a predictive coding inspired architecture can lead to BOS signals, with properties very similar to those in the brain, even without explicitly training the network to localize or identify objects in visual scenes.

A prior study showed that PredNet units signal illusory contours and end-stopping^17^. The emergence of BOS signals as well as these other extra-cRF phenomena under the predictive coding framework raises a question: do these phenomena result from a single hierarchical neural computation? Several lines of prior research are consistent with that possibility^17,30,33,34,36,48^. A complete answer to this question is hard to obtain by solely doing physiology experiments: detailed maps of neural connections are often unavailable, and it is challenging to precisely manipulate these connections. ANNs have the unique advantage of possessing complete connection profiles^22,49,50^, and allow one to perform ablation studies. Our work thus establishes PredNet as a useful complementary tool towards achieving an understanding these computations.

PredNet’s E modules have been interpreted as being akin to superficial layers (L1/2/3), and the R modules as akin to deeper layers (L5/6) of the visual cortex, following the proposed functional specialization of cortical layers in predictive coding^17,27,33^. The presence of BOS signals in both E and R modules aligns with physiology, where BOS neurons exist in both superficial and deeper layers^3^. However, one should be cautious to equate E and R modules to different cortical layers. For example, the R modules have lateral connections, which the E modules lack these, unlike in physiology where lateral connections exist in both superficial and deep compartments^36,51,52^. Further studies are needed to understand the functional role of different areas and layers in this hierarchical computation and the communication between them. Because of the flexibility to manipulate network architecture and connections, ANNs are a useful complementary tool in such studies^17,27^.

Our discovery of BOS units in PredNet and the ablation experiments indicate that BOS neurons may be useful for video prediction. To predict future visual input, it is useful to predict object motion^53^. Objects typically move as a whole, i.e. pixels within object boundaries most likely move together^54^. Because BOS units indicate which pixels belong to an object surface, they may help to predict by allowing the system to easily apply a uniform optical flow to objects. Indeed, in computer science, incorporating optical flow^55,56^, disentangling object motion from content^57–60^, and separating foreground objects from background^54,61^ have been shown to improve video prediction performance. Beyond video prediction, in object recognition, deep neural networks have been criticized for relying mostly on textural information to recognize object categories rather than on object shapes^62–64^, in contrast to human visual perception^65,66^ (but perhaps more akin to mouse visual perception^67^). Explicitly embedding a BOS unit module may guide neural networks to rely more on shapes, and potentially achieve more robust recognition as well as prediction.

Overall, our work demonstrates that brain-like BOS signals emerge in a self-supervised network trained to predict future input. This implies a shift from the traditional view of BOS as a static ‘what stream’ operation towards a computation that is highly beneficial to predict future input in natural dynamic environments.

## Methods

### PredNet architecture

In this study, we utilized the artificial neural network PredNet, which was developed and trained by Lotter et al. (2017)^26^ (code is available at: https://github.com/coxlab/prednet). Here we briefly summarize PredNet’s architecture and how it was trained. PredNet is an artificial neural network (ANN) that has four layers (labeled as ‘*l*’). Each layer consists of four types of modules: the Representation module (*R*_*l*_), the Prediction module (Â_*l*_), the Prediction target module (*A*_*l*_), and the Prediction Error module (*E*_*l*_). Updating unit activities in PredNet involves two main stages at each time step:

Top-to-Bottom Update: The network updates the R modules from top to bottom at each time step. Each *R*_*l*_ module gets inputs from the *R*_*l*+1_ module and the *E*_*l*_ module. This updating process goes from the *R*_3_ module to the *R*_0_ module in sequence. The *R*_0_ module then generates a predicted current video frame (Â_0_). Bottom-to-Top Update and Error Calculation: The update process then reverses, proceeding from bottom to top. The network calculates the prediction error by comparing Â_0_ with the actual next video frame, *A*_0_. This error is bifurcated into positive and negative parts (akin to biological ON-center and OFF-center neurons). Positive and negative errors are grouped in the *E*_0_ module. *E*_0_ then outputs a target prediction *A*_1_, which gets compared with Â_1_ produced from *R*_1_. The error from this comparison is the *E*_1_ module. The network continues this process up to the final layer (layer 3).

Mathematically, the PredNet dynamics are defined by

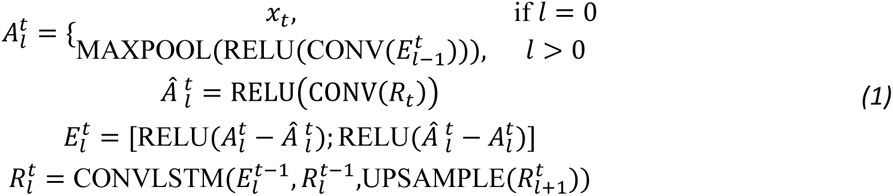

where *t* is the time step, *x*_*t*_ is the actual video frame. ConvLSTM uses a tanh activation function, which means that the R module activation can be negative (possible values range from -1 to 1). Because biological neurons do not have negative spike rate, PredNet unit’s response was defined in this study as the unit activation plus one, i.e. the response baseline was shifted by +1 in all modules (after PredNet’s computation was completed, thus this did not affect PredNet’s algorithm). The PredNet architecture contains 3, 48, 96, and 192 convolution channels in layers 0 to 3, respectively. The input image size is 128 by 160 pixels. The number of units in R modules are respectively 61,440 in *R*_0_, 245,760 in *R*_1_, 122,880 in *R*_2_, and 61,440 in *R*_3_. The *A*_*l*_ and Â_1_ modules have the same number of units as the *R*_*l*_ module. Due to the bifurcation of positive and negative error, E modules have twice the number of units compared to the R modules.

The training loss function is applied on the prediction error

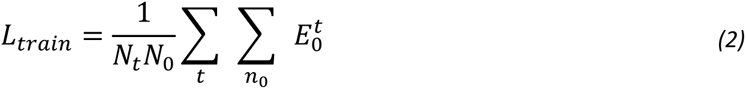

where *N*_*t*_ is the number of time steps used in training, *N*_0_ is the number of *E*_0_ units. The training utilized the KITTI dataset, which contains videos recorded from car-mounted cameras in Germany. Videos were segmented into sequences of 10 continuous frames. These frames were then center-cropped and downscaled to a resolution of 128 by 160 pixels. The parameters of PredNet were optimized using backpropagation with the Adam optimizer.

### Classical Receptive Field of Units

We measured the classical receptive field (cRF) of units in PredNet using sparse noise stimuli (SI Figure 1), similar to the approach used in physiology. We created an image (128 x 160 pixels) with one pixel set to either white or black, while all others were set to gray (gray level = 0.5, scales from 0 to 1). 40,960 (128 × 160 × 2 where the factor two is for black and white pixel) unique images (128 x 160) were generated, each featuring a distinct single pixel, in either white or black. These images were repeated for four time steps, yielding a total of 40,960 sequences. For each unit, recorded activity to these sequences was summarized into two heatmaps (each size 128 × 160), each representing responses to respectively white-pixel and black-pixel scenes. For example, the white heatmap’s *i*, *j* entry is the single unit’s time-averaged response to a scene with a white pixel located at *i*, *j* (gray otherwise).

The two heatmaps (for one unit) were then z-scored and converted to absolute values. These heatmaps were merged into one heatmap by taking the maximum absolute values for each entry. This merged heatmap summarizes the unit’s maximum response to a pixel at each location irrespective of its color (white or black). The cRF for each unit was defined as the union of the pixel positions for which the absolute values of the maximal z-scores across both heatmaps exceed 1.

### Standard Square Stimuli

Scenes with square objects are commonly used in neurophysiological studies to assess whether a unit is selective for BOS^1–3,23^ and this selectivity is known to extend to natural images^5^. We used similar scenes, consisting of a square with a size (width) of 50 pixels, positioned with one border centered at the center of the scene (central border). The color of the square and the background can be either light (gray level = 0.33 on a scale from 0 to 1) or dark gray (gray level = 0.66), but they are always different from each other in a given scene. These square scenes can be defined mathematically by three parameters. The first parameter, α, denotes the square’s orientation, with a range from 0 to 180 degrees. The second parameter, γ, is a binary variable indicating which side the square is given a fixed orientation (i.e. side of ownership). The final parameter, γ, is a binary variable that indicates the contrast polarity across the central border. For each square orientation defined by α, there are four possible square scenes, determined by different combinations of and γ. Each of these scenes is repeated over 20 time steps. We used 10 different orientations (equally spaced by 18°).

### Candidate Unit Selection

To define selectivity for border ownership, it is important to verify that the units under examination respond to changes in border ownership rather than to low-level stimulus changes within the cRF. Therefore, similar as in neurophysiology studies, we restricted our analysis to units that passed the following two criteria (termed ‘candidate units’). First, the unit’s cRF must include the center of the scene. Because the central border of the square scenes was placed exactly in the scene center, this ensured that the unit’s cRF includes the center of this border. Second, the cRF must fit within a circle centered at the center of the scene and with a radius of 20 pixels. Because the square size (width) is 50 pixels, this makes sure that the cRF does not overlap with any other border of the square besides the central border.

### Averaged Border Ownership Index across orientations (*B*_*av*_)

Similar to neurophysiology studies^1–3^ we quantified tuning for border ownership using the Border Ownership Index (BOI). This is computed from the response of PredNet units to standard square scenes.

The BOI is defined as

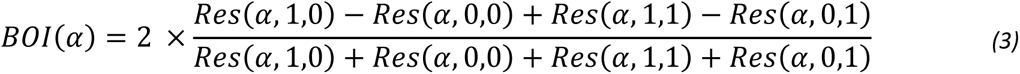

where *RES*(α, β, γ) is the unit’s time-averaged (between 0 and 19 time steps) responses to a square scene specified by orientation α, side-of-ownership γ and contrast polarity γ. The sign of the BOI thus indicates which side (γ) of BOS (for a given orientation) the unit prefers, and the magnitude indicates the strength of the BOS tuning.

To evaluate the overall BOS selectivity across orientations, we defined *B*_*av*_ as the circular average of the BOI across α. Similar to BOI, the magnitude of *B*_*av*_ is a measure of the strength of BOS tuning, and its angle indicates the unit’s preferred side of BO.

We evaluated the statistical significance of *B*_*av*_ using a permutation test. In this test, we shuffled the labels that signified the side of BOS (γ) for each orientation α. These data were then used to compute a shuffled BOI(α) and *B*_*av*_. This procedure was repeated 5,000 times to generate a set of 5,000 *B*_*av*_ values after shuffling, for each unit. Denoting the quantile of the unshuffled *B*_*av*_ among the shuffled *B*_*av*_ as Q, the p-value (two-tailed) was estimated as 2 × *min*{*Q*, 1 − *Q*}. Units with a p-value less than 0.05 were defined as BOS units. 95% confidence intervals on proportions of units for which *B*_*av*_ was significant were computed using Wilson score^68^.

Note that the values of *B*_*av*_ and BOI reported here cannot easily be compared with similar indices in neurophysiology, because these values change when the DC level of unit activity is changed. As mentioned above, to avoid negative values for unit activity in PredNet, we arbitrarily increased activity levels by +1. Furthermore, the average BOI across time depends on when the response starts relative to the duration of the analysis window. This is at ∼50% of the window duration for the unit shown in Fig. 1D (top), whereas in physiology studies this is typically closer to ∼10%. For example, the activity functions shown in Figure 1D (top panel) show a BOI of 0.0149 at time step 10, but computing this without adjusting the unit activation (i.e. without +1) leads to BOI = 0.68. Zhou et al. use ‘response ratio’ to quantify the magnitude of BOS tuning, defined as the ratio of the mean response to non-preferred BOS over the mean response to preferred BOS. For the activity functions shown in Fig. 1D (top panel) this value is 0.561(averaged across analysis window), well within the range of values found for neurons in the macaque visual cortex^1^.

### Analysis of BOS Unit Responses to Different Square Orientations, Positions, and Sizes

In these experiments, varied parameters were square orientation (α), side-of-ownership (γ), contrast polarity (γ), position along the orientation (*d*), and size (*s*). We first measured the response to a set of four standard square scenes (Figure 1A). For each unit, the orientation α is fixed at the orientation with the maximum absolute BOI. The position is zero, indicating that the square border intersects exactly with the scene center, and the square size (width) is 50 pixels. BOS units’ responses were averaged over time and contrast polarity. The γ value with the larger averaged unit response was defined as the preferred side (γ_*p*_), whereas the opposite was defined as the non-preferred side (γ_*p*_). These preferences were solely determined by the standard square scenes.

We then examined the effect of changing square size. All other parameters remained the same as in the standard square scenes stated above, except for square size. Eight square sizes were used, ranging from 10 to 80 pixels. For each unit *i* and each square size *s*_*j*_, we computed the responses averaged across time and contrast polarity, yielding 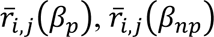. We then normalized two response arrays of each unit 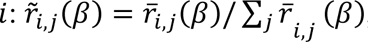, where γ can be γ_*p*_ or γ_*np*_ Figure 2D (left panel) displays the time course of 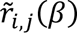 across units *i*. For each unit *i* and square size *s*_*j*_, we computed a BOI as the difference in response between the γ_*p*_ and γ_*np*_, i.e. 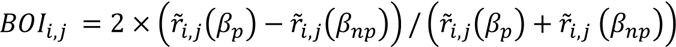. We performed a bootstrapping test to assess statistical significance of this metric. A BOI dataset consisted of *D^*B*OI^* = {*BOI*_*i,j*_ } for all units *i* and square sizes *s*_*j*_. We obtained 10,000 bootstrap samples ^*s*^ from this dataset. For each *D*^*s*^, we computed an averaged BOI, denoted as *BOI*^*s*^. The p-value was estimated as *p* = 1 – *Q*, where *Q* is defined as the quantile of 0 among all *BOI*^*s*^. If p-value was smaller than 0.05, we concluded that the BOI averaged across size was statistically significantly positive in the population of BOS units (SI Figure 4).

The same procedures apply to varying square position, simply replacing square size with square position (SI Figure 4). Fifteen square positions were used, ranging from -30 to 26 pixels.

When examining the unit’s response to different orientations, we created square scenes with 10 possible orientations (equally spaced between 0 and 180 degrees), keeping the position at 0 and size at 50 pixels. Units’ responses were collected to compute the BOI for each orientation using equation (3). Data from one example unit is shown in Figure 2A (left panel). To evaluate the statistical significance of BOI for a given orientation, we compared the unshuffled BOI to that in a null distribution. Unlike biological neurons, which differ in response from trial to trial, PredNet does not have noise. In order to obtain sufficient data to generate a shuffled distribution, for each orientation, we varied square size (10 different sizes were considered, conceptually mimicking 10 “repeated trials”). The unshuffled BOI for a given orientation was computed for this orientation across square size. The null distribution for BOI distribution was obtained by shuffling the labels indicating the side-of-ownership γ (i.e., border ownership), separately for each square size and contrast polarity (5,000 shuffles). The quantile (*Q*) of the unshuffled BOI within shuffled BOI set was computed. The p-value (two-tailed) was estimated as as 2 × *mi*{*Q*, 1 − *Q*}. If the p-value is less than 0.05, BOI along an orientation was said to be statistically significant (indicated in Fig. 2A as filled circles). The above procedure resulted a subset of orientations with statistically significant BOIs. The span of each BOS unit was computed as the difference between the two most distant preferred object locations (circular distance between the two angles corresponding to those locations). 95% confidence intervals on proportions of units for the span was smaller than a certain value were computed using Wilson score^68^.

### Square Fragment Stimuli

The squares in the square scenes can be divided into eight fragments^31^: the Central Edge (CE), which is the one in the middle of the scene; there are two Near Corners (NC), two Near Edges (NE), two Far Corners (FC), and one Far Edge (FE). To examine how thee fragments modulate the activity of BOS units, the four standard square scenes (Figure 1A, orientation aligns with the preferred BOS orientation for each unit) were converted into fragmented square scenes, as described below.

To isolate one fragment, a 2D Gaussian filter (σ = 5 pixels) was applied at the fragment’s center. This kept the fragment’s central region largely unaltered, while the parts of the scene further away gradually fade to a uniform gray (gray level = 0.5 on a scale from 0 to 1). For scenes with multiple fragments (e.g. ‘All’), a Gaussian filter was applied to each fragment. Note that the smallest distance between two fragment centers is 25 pixels, thus much larger than σ, resulting in negligible interference between filtered fragments at different locations.

Using this Gaussian filter method, we created 9 scenes with a Central Edge (‘with-CE’) and 9 scenes without a Central Edge (‘without-CE’) for each of the four standard square stimuli (Figure 1A). Among the with-CE scenes, one scene only has the CE fragment, seven scenes have the CE and one additional fragment, and one scene has all fragments. The without-CE scenes are similar to the with-CE scenes, except that they do not contain the CE fragment. Thus the “all fragments” without-CE scene contains 7 fragments. Each scene is presented during 20 time steps.

### Processing of Units’ Responses to the Square Fragment Stimuli

The NC fragment could potentially partially intersect with the cRF. To prevent this, we limited this analysis to the subset of BOS units whose cRFs fitted within a circle of 30 pixels diameter centered at the center of the scene. This more conservative selection yielded 30, 145, 71, 5 units from respectively E0 to E3, and 1, 3, 2, 0 units from respectively R0 to R3. For with-CE scenes, the surround influence of square fragment X is defined as the unit’s response to the X + CE scene subtracted by the response to the CE scene. Similarly, for without-CE scenes, the surround influence of square fragment X is defined as response to the X scene subtracted by a full gray scene. If X is FE, the surround influence of X is computed as above. Otherwise (X = FC, NE, NC), the surround influence of X is the average of the surround influences of two conjugate edges (e.g., CE1 and CE2).

The surround influences of X for all BOS units were computed, resulting in a list where the length equals the number of BOS units. To avoid bias in mean estimation due to outliers, outliers (1.5 interquartile range below the first quantile or above third quantile) were removed before computing the sample mean and SEM (Fig. 2F and SI Fig. 5). However, all units were included when performing statistical tests (indicated by figure caption).

### Square-Ambiguous, Square-Opposite Square, and Figure-Off Sequences

Each trial in the Square-Ambiguous sequences consisted of 20 time steps, broken down into two phases. Initially, Scene 0, one of the four standard square scenes (Figure 1A), was displayed during four time steps (T0 = 4). Subsequently, Scene 1 was shown during 16 time steps (T1 = 16). Scene 1 only contained a central border that divides the whole image into a left and a right half; hence the side of ownership of this border was ambiguous. The contrast polarity and orientation of Scene 1 were consistent with Scene 0 (i.e. the information in the cRF was the same).

Similarly, the Square-Opposite Square sequences started with one of the four standard square scenes as Scene 0. Scene 1 was a version of the square scene with reversed BOS, but maintaining contrast polarity for the central border. For example, if Scene 0 was panel 1 in Figure 1A, then Scene 1 was panel 2 in Figure 1A.

For Figure-Off Sequences, Scene 1 was always a full gray. Scene 0 depended on the subtypes: Ambiguous-Off, Grating-Off, and Pixel-Off sequences. For Ambiguous-off, Scene 0 was an ambiguous border. It had two versions that vary in contrast polarity. In Grating-Off sequences, Scene 0 was a grating with a 10-pixel spatial period, and it had two versions with grating phases of either 0 or 180 degrees. For Pixel-Off sequences, Scene 0 was gray except for a single pixel at the center, which was either white or black corresponding to two versions.

All scenes were generated such that the orientation corresponds to that for which each unit’s |BOI| was maximal.

### Relative Response Difference

The Relative Response Difference (RRD, used in the result section “PredNet’s BOS units exhibit hysteresis, similar to BOS neurons in the brain” and Figure 3) is (*a* − *b*)/(*a* + *b*), where *a* indicates the time-averaged response to preferred stimuli, and *b* indicates the time-averaged response to non-preferred stimuli. Which stimulus was preferred only depended on the averaged response to Scene 0.

RRD half-life was defined as the earliest time after the scene switch where the absolute value of RRD was less than half of its maximum. The half-life across the three types of Figure-Off sequences were averaged in Figure 3D. For this analysis, we only included units for which the half-life of all three types of Figure-Off sequences could be measured (exclude RRD that never dropped to half of its maximum within the analysis window). This yielded 10 out of 22, 8 out of 25, 9 out of 12 and 1 out of 2 BOS units in respectively R0, R1, R2 and R3; and 25 out of 32, 160 out of 199, 90 out of 131, and 15 out of 22 BOS units in respectively E0, E1, E2 and E3. The Wilcoxon signed-rank test was used to compare half-life between Square-Ambiguous sequences and Figure-Off sequences.

### Three video types for Ablation Experiment

We generated three types of videos to evaluate PredNet’s prediction performance (examples shown in SI Figure 7). (1) Translating Square videos include a square that moves at a constant speed and direction. Square size is 50 pixels and oriented such that the central border had a vertical orientation (square gray level = 0.33 and background gray level = 0.66 on a scale from 0 to 1). The initial position and velocity of the square were chosen such that the square was always in the scene center in the 10^th^ frame. Forty translating square videos were created, corresponding to 40 evenly spaced moving directions (equally spaced between 0 and 360 degrees). (2) Random Square videos: each of these videos featured a random number of squares (between 1 and 5). At the beginning of each video, each square’s central position was randomly set in the scene. The size of each square was also randomly chosen (between 10 and 50 pixels), and the x and y components of each square’s velocity were randomly set at a value between -2 and 2 pixels/frame. Forty random videos were generated. (3) KITTI testing videos: 41 videos from car-mounted cameras were used, which were not used during PredNet’s training^26^. For all video types, each video consisted of 20 frames.

### Subsampling BOS and Non-BOS Units to Reduce Their Response Differences

Unit activity in response to the videos were squared and averaged across all videos and time steps for each video type, resulting in Mean Squared Response (MSR). For each module and video type, we have two sets of MSR, one for the BOS units and another for the non-BOS units, denoted as 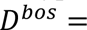 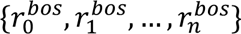 and 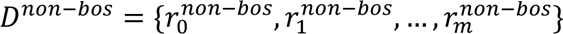, respectively, where *n* and *m* representing the number of BOS and non-BOS units in one module.

For each of the *D*^*bos*^ and *D*^*non*−*bos*^, we subsampled *k* = *min*, {*n, m*} units (1,000 samples). This resulted in 1,000 pairs of sampled datasets, denoted as 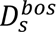, and 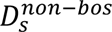 with *s* ranging from 1 to 1,000. For each pair, we computed a score to measure the similarity between datasets in a pair

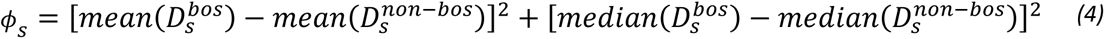

where the *mean*(⋅) and *median*(⋅) represent those quantities of the dataset. The dataset pair with smallest score ϕ_s_ was subjected for further statistical analysis, using the Wilcoxon rank-sum test and the t-test. If both p values were larger than 0.5, we considered the dataset pair as our final subsampled datasets. If not, we reduced *k* by 1 and repeated the procedure above. This whole procedure makes sure that both BOS and non-BOS populations have the same number of units (equal to *k*), and their MSRs do not show significant difference. SI Figure 8 displays the MSR of the obtained subsampled unit populations.

### Compute the Prediction Error of the Ablation Experiment

For each video type, we created *N*_α_ bootstrapped samples, each containing *N*_*a*_ videos. We denoted 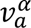 as the *a*^*t*ℎ^video in the α^*t*ℎ^ bootstrapped sample, with α ranging from 0 to *N*_α_ − 1, and *a* from 0 to *N*_*a*_ − 1. In this study, *N*_α_ = *N*_*a*_ = 10.

For each video 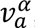, we performed the ablation experiment several times, for different samples of ablated units, in each module separately. We varied the number of ablated units *n* (ranges from 1 to *min*_*bo*_, *N*_*non*−*bo*_}, where *N*_*bo*_ and *N*_*non*−*bo*_ indicate respectively the number of BOS and non-BOS units available in the module). For each *n*, we generated *N*_*u*_ = 10 bootstrapped unit samples from the unit pool (i.e. either from the BOS/non-BOS unit population in each module). A single sample is denoted as 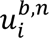, where *b* is a Boolean variable indicating whether the ablated units are BOS units or non-BOS units, and *i* = 1, 2, …, *N*_*u*_ represents the *i*^*t*ℎ^ unit sample. For each ablation sample 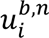, the unit activity in the sample was set to zero. Mean-squared prediction error (MSE) was measured as the mean-squared difference between the predicted (Â_0_) and actual frames (*A*_0_), averaging over all pixels and time steps. Relative prediction error (RPE) of one video and one ablation sample was computed as

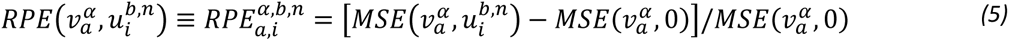

where 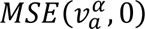 represents the MSE to the same video without ablation. We then computed the average RPE for a single video sample α and a given number of *n* ablated units:

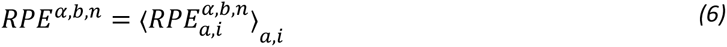

where ⟨⋅⟩*_a,i_*, represents the average across indices *a* and *i*. Dots and error bars in Figure 4 show the mean and SEM of *RPE*^α,*b*,*n*^ across different video samples α, with respect to the number of ablated units *n*, for the subsampled population (see previous Methods section: ‘Subsampling BOS and Non-BOS Units to Reduce Their Response Differences’). SI Figure 9 shows the result for the original population (without subsampling).

### Statistical Analysis of the Ablation Experiment

We model the *RPE*^α,*b*,*n*^ as a linear model

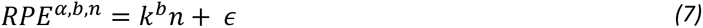

where the intercept term is zero because the RPE is zero when no units are ablated. ε is an error term with a zero mean and a constant unknown variance, and *k*^*b*^ is the slope of a line that represents the average change in RPE if one additional unit is ablated (*b* = *bos* for ablation of BOS units, *b* = *non* − *bos* for ablation of non-BOS units). We are interested in determining whether the slope *k*^*bos*^ is significantly different from *k*^*non*−*bos*^. A bootstrap method is used as follows.

Observations are denoted as *D*^*b*^ = {*RPE^α,b,n^*} where α and *n* indicate respectively video samples and number of ablated units. *N*_*s*_ = 10,000 bootstrap samples are generated by resampling *D*^*b*^ with replacement, denoted as *D*^*b*,^ where *s* = 1,2, …, *N*_*s*_. For each bootstrapped dataset, we used ordinary least squares linear regression to compute a slope *k*^*b*,^. 95% confidence interval of the slopes were estimated from the bootstrapped distribution (shown as error bands in Fig. 4 and SI Fig. 9). Subtracting the two slope sets, we got *N*_*s*_ slope differences denoted as Δ*k*^*s*^ = *k*^*bos*,*s*^ − *k*^*non*−*bos*,*s*^. The p-value (two-tailed) was then estimated as 2 × *min*{ *Q*(0, {Δ*k*^*s*^}), 1 − *Q*(0, {Δ*k*^*s*^}) } where *Q*(0, {Δ*k*^*s*^}) is the quantile of 0 in the set of slope differences {Δ*k*^*s*^}.

## Funding

NIH grant R00EY031795 (TPF)

Incubator for Transdisciplinary Futures: Toward a Synergy Between Artificial Intelligence and Neuroscience (RW).

## Author contributions

Conceptualization: ZY, RW, TPF

Methodology: ZY, RW, TPF

Investigation: ZY

Supervision: RW, TPF

Writing: ZY, RW, TPF

## Data and materials availability

## Supplementary Materials

**SI Figure 1.**
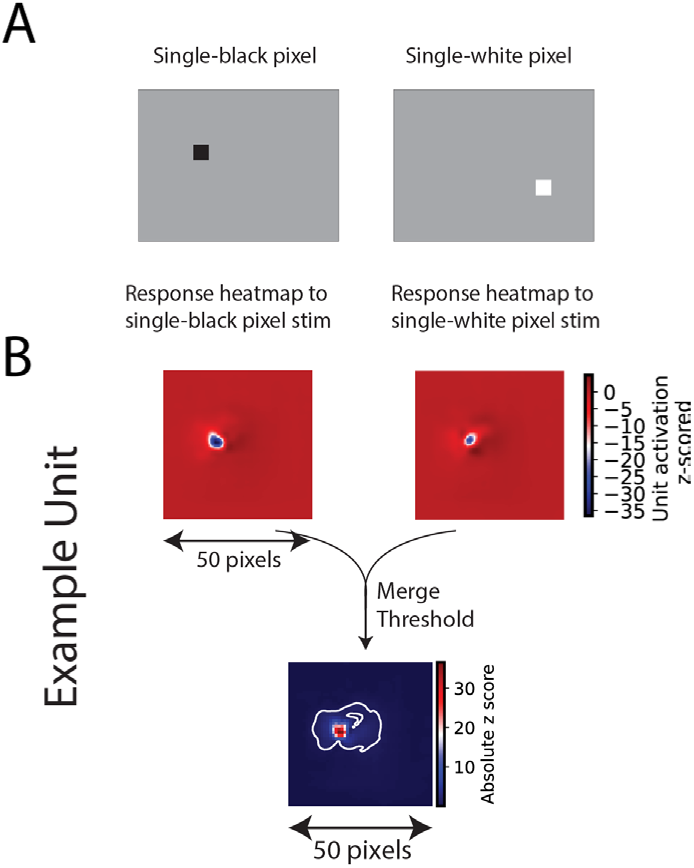
Illustration of the method to measure the cRF of PredNet units. **(A)** A sparse noise scene is a gray scene with only one black or white pixel, at a random position. These scenes were used as input to PredNet over four time steps. **(B)** cRF for an example unit. The unit’s responses to the sparse noise scenes were collected and normalized (z-scored) into two heat maps, one for black pixel noise and the other for white. Each value in the black or white heatmap corresponds to the unit’s normalized response to a black or white pixel at the same entry position. The two heatmaps (for one unit) were merged into one heatmap by taking the maximum absolute values for each entry. Positions with an absolute value of the z-score greater than 1 were defined as the cRF (indicated by white contours).

**SI Figure 2.**
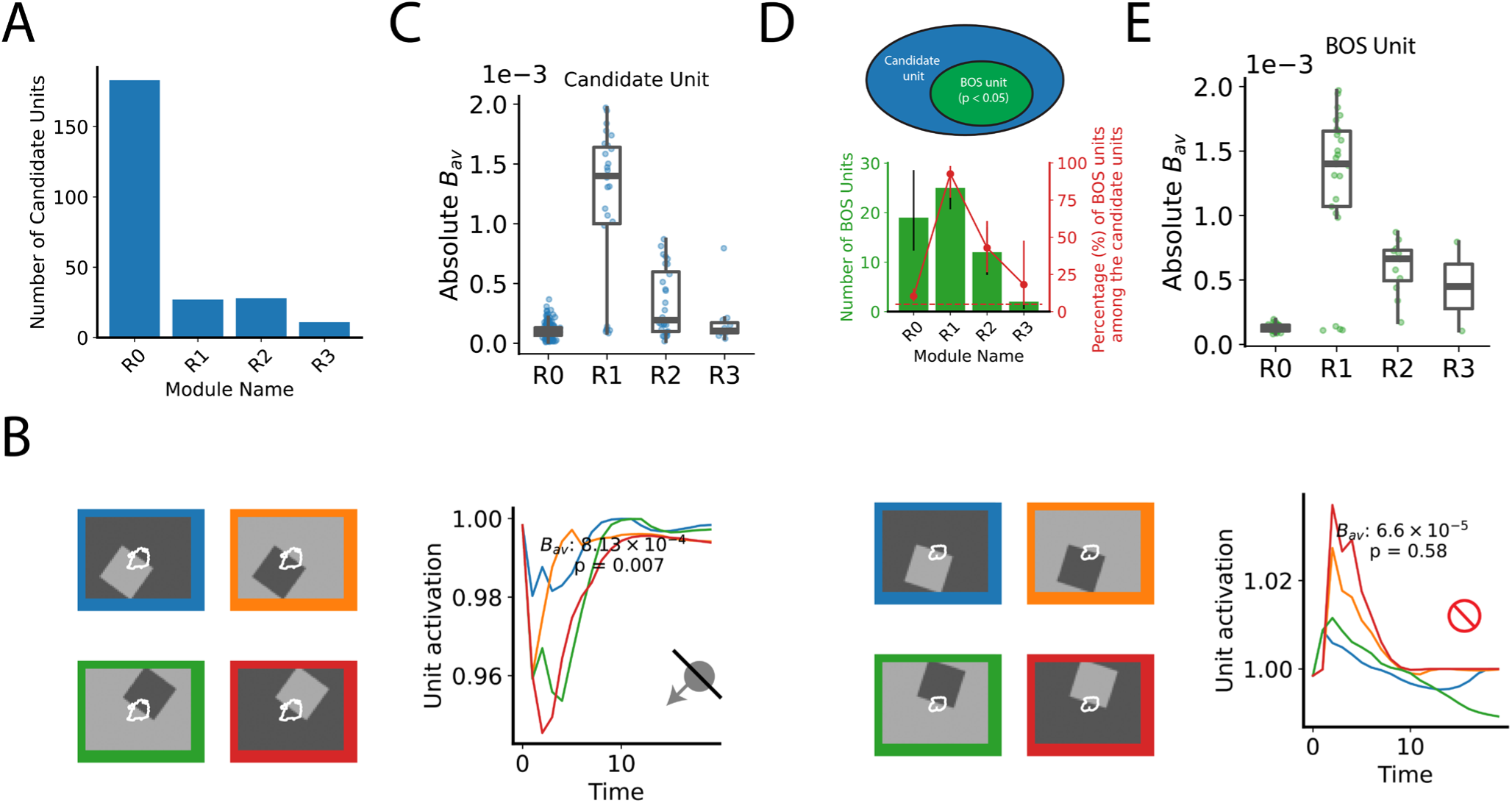
BOS units emerge in PredNet’s R modules. (**A**) The number of candidate units in R modules across different layers. (**B**) Responses of two example units (module *R*_2_), with white contours indicating the cRF (similar to Fig. 1D). *B*_*av*_ measures the unit’s response different to different BOSs across different square orientations (see Methods). P value (two tailed) was computed by comparing *B*_*av*_ to that after shuffling stimulus labels (permutation test, see Methods). Arrow in the middle-left panel indicates the preferred side of BOS for the example candidate unit. (**C**) The *B*_*av*_ distribution of the candidate units in different R modules. Each dot is one candidate unit. (**D**) Among the candidate units, units with p-value smaller than 0.05 are defined as BOS units. Error bars indicate 95% confidence intervals. Horizontal dashed line indicates chance level of 5%. (**E**) The *B*_*av*_ distribution of BOS units in different R modules. Each dot is one BOS unit.

**SI Figure 3.**
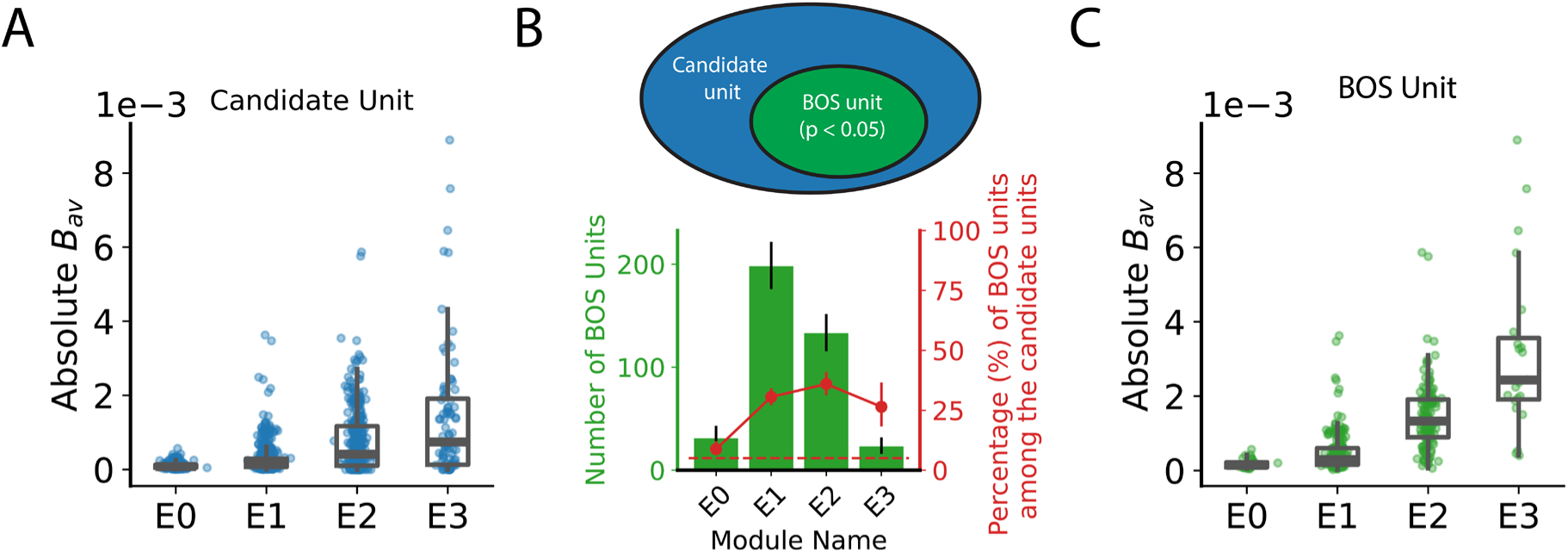
*B*_*av*_ distribution of units in E modules. Similar as SI Fig. 2C-E, for E modules.

**SI Figure 4.**
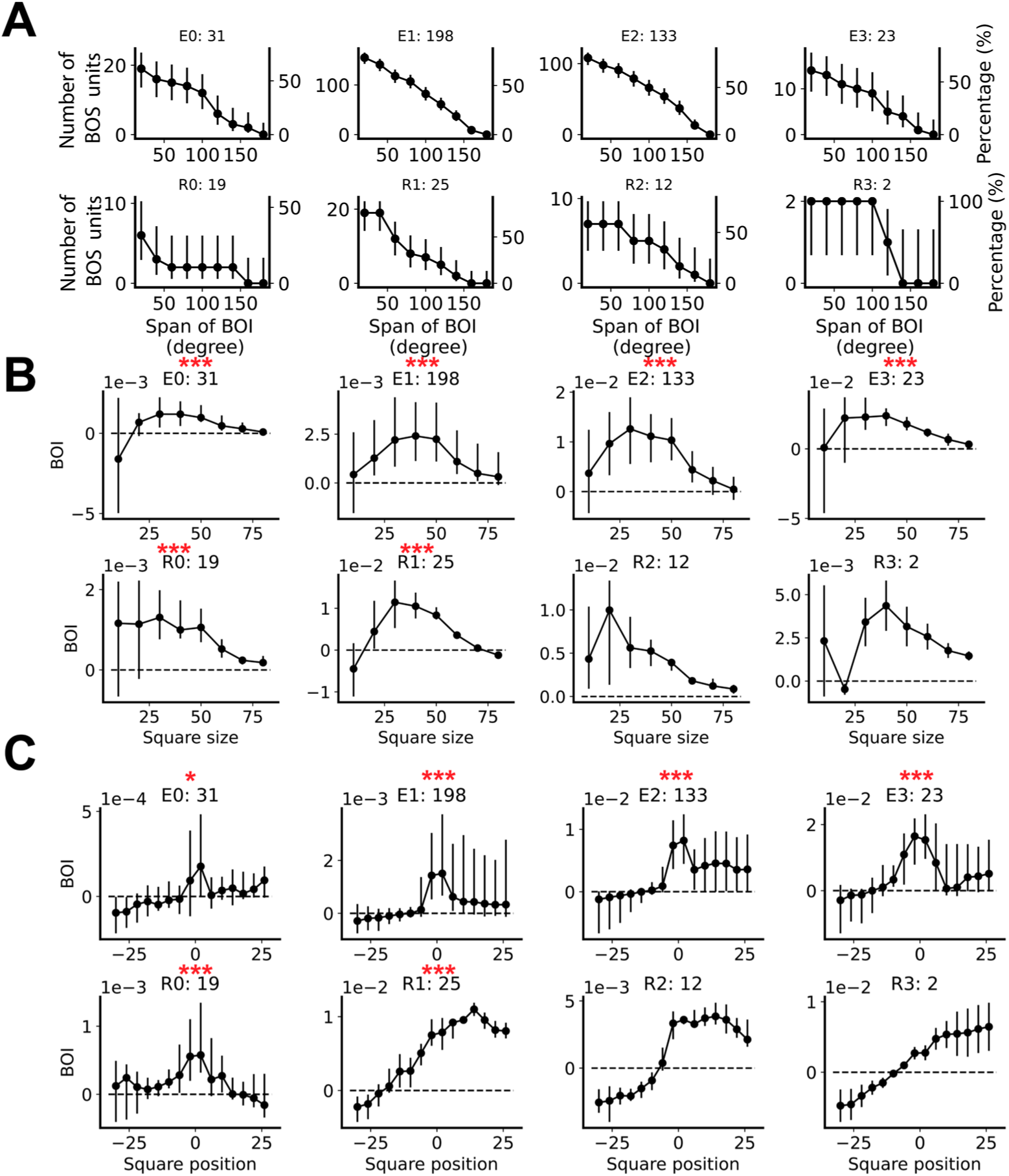
BOS signals are robust to different stimulus parameters. (**A**) Similar to Figure 2B, for other modules. (**B, C**) BOI across different square sizes and positions. The dots and error bars represent the median, first and third quartiles across all units in a module. The number after the module name in the panel titles denotes the total number of BOS units included per module. Red symbols indicate whether the averaged BOI across conditions (square sizes or positions) are statistically significantly larger than zero, ***: p < 0.001; *: p < 0.05; bootstrapping test (see Methods). Statistical significance was only evaluated in modules with more than 15 BOS units.

**SI Figure 5.**
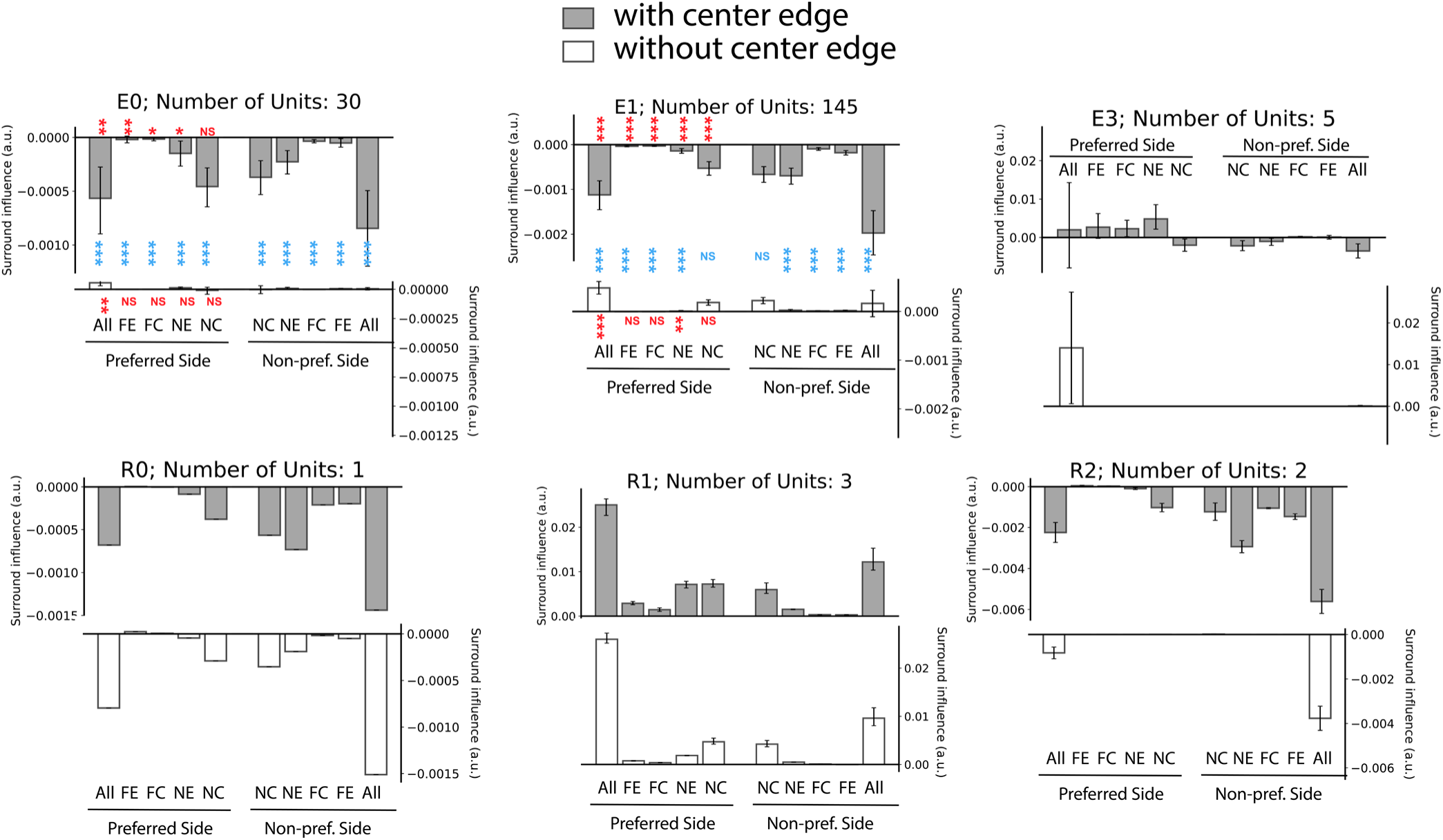
BOS units’ responses to square fragments on the preferred side of BOS are generally larger than on the non-preferred side of BOS. Similar to Fig. 2F, for BOS units in different PredNet modules. Red text indicates whether the surround influence for a particular condition is significantly larger on the preferred side than on the non-preferred side. Blue text indicates whether the absolute value of surround influence of with-CE is significantly larger than without-CE case. Wilcoxon signed-rank test. ***: p < 0.001; **: p < 0.01; *: p < 0.05; NS: no significance.

**SI Figure 6.**
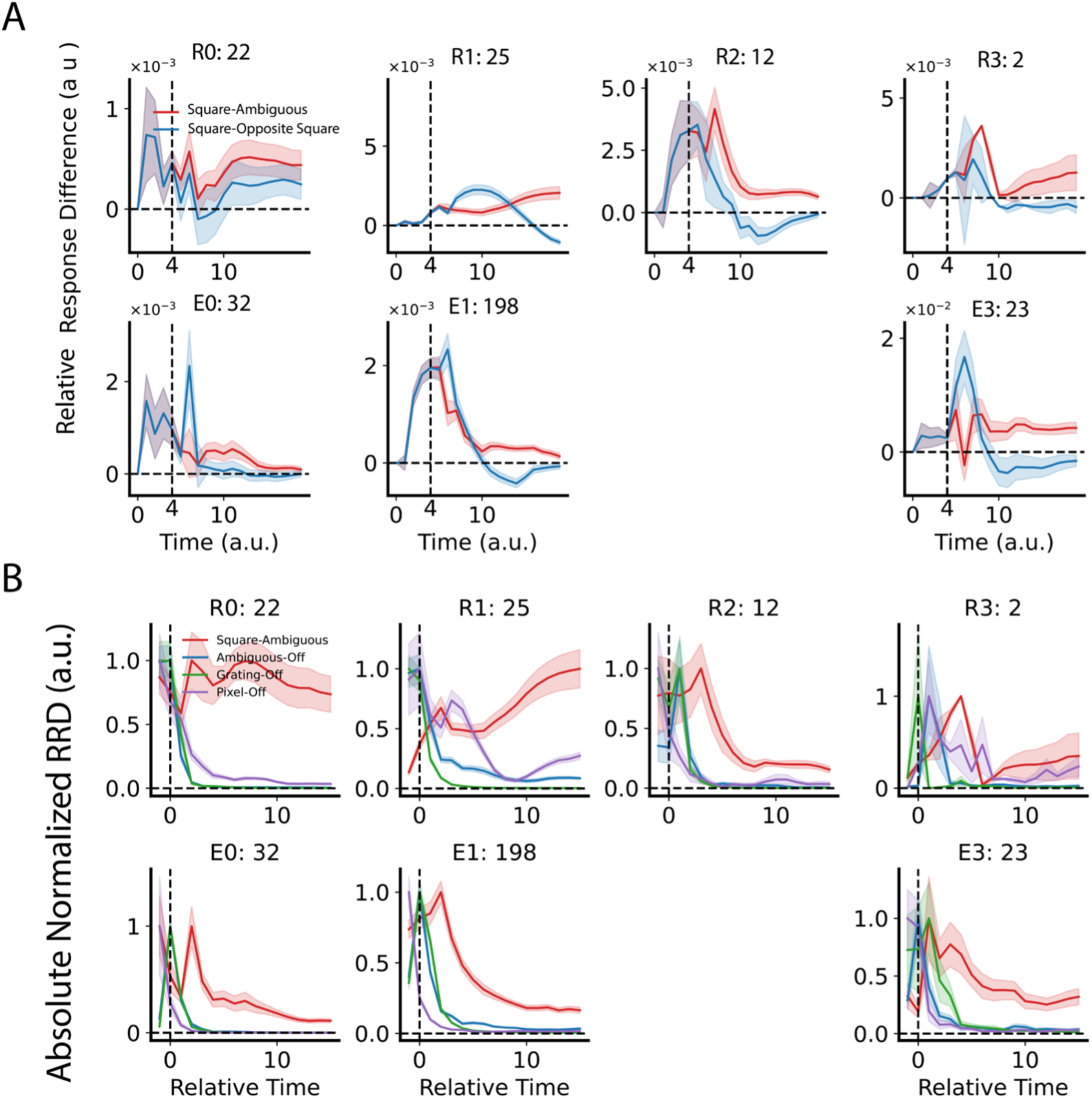
Persistent BOS signals in different modules. (**A**) Similar as Fig. 3B, for other modules. The number of BOS units in each module is indicated in the title. (**B)** Similar as Fig. 3C, for other modules.

**SI Figure 7.**
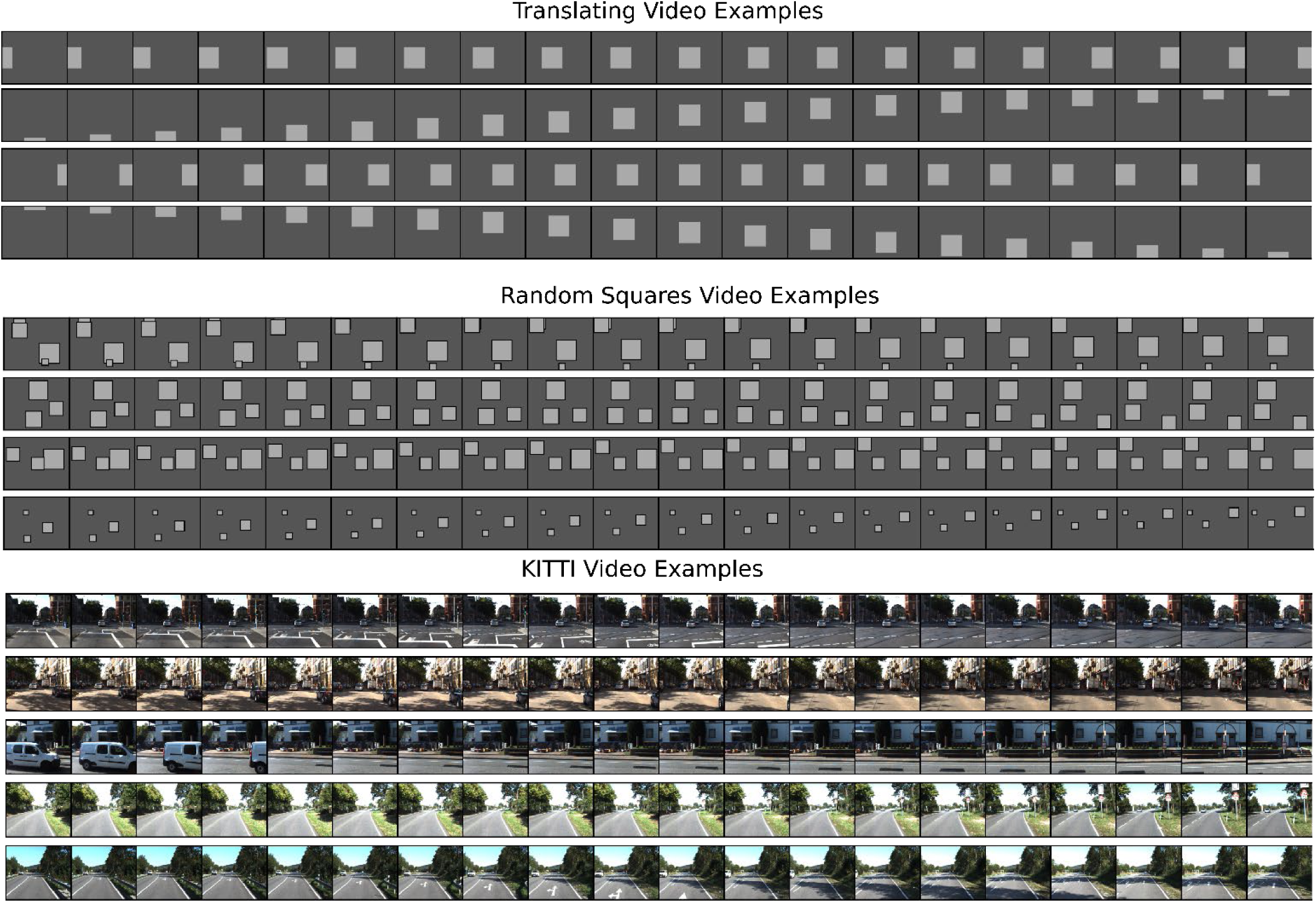
Three video types used for the ablation experiment. Figure shows example videos from each video type. Each row shows a different unique video for each of the three types. Video length is 20 frames, shown during 20 time steps.

**SI Figure 8.**
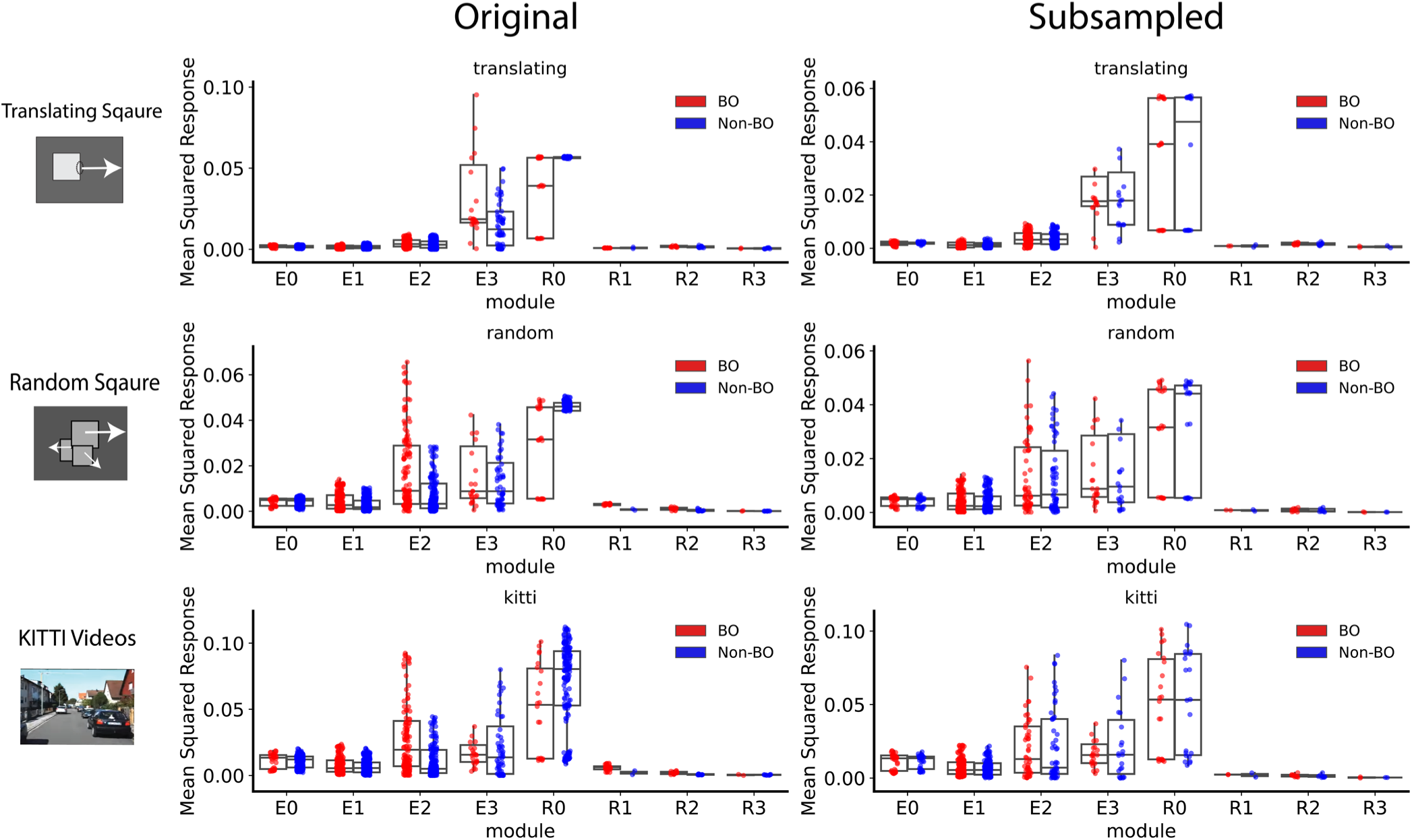
Activity in subsampled BOS and non-BOS unit populations and original populations. For each unit, mean squared response is the square of the averaged response, averaging cross time and videos. Each dot is one unit’s mean squared response. Boxes indicate the interquartile range between the first and third quartiles with central mark inside each box indicating the median. Whiskers extend to the lowest and highest values within 1.5 times the interquartile range. Outlier units not shown for better visualization (but included in the metrics indicated by the boxplots).

**SI Figure 9.**
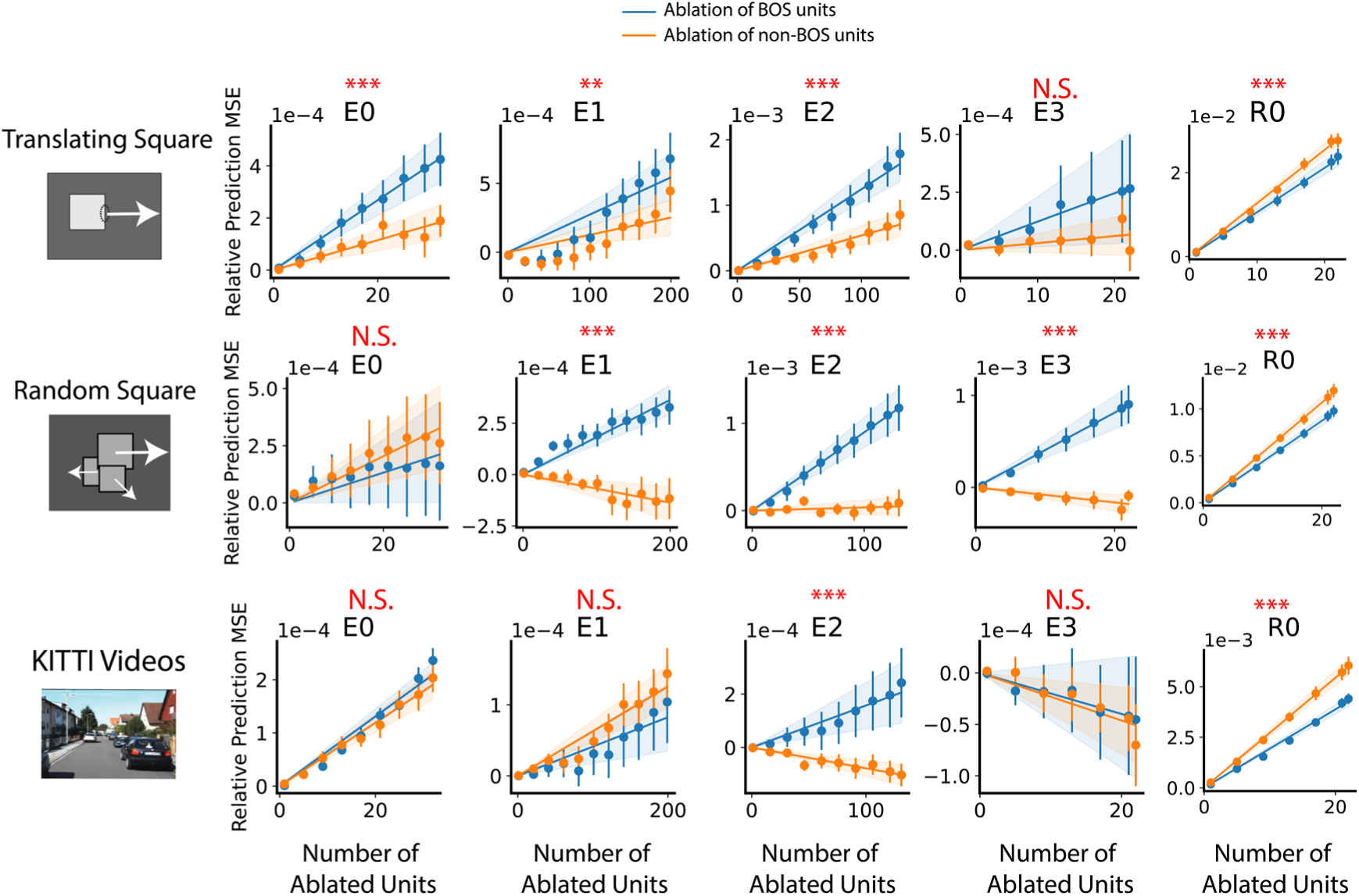
The effect of ablating the original BOS/non-BOS units, without subsampling. Similar as Figure 4 but using original unit population (no subsampling).

**SI Figure 10.**
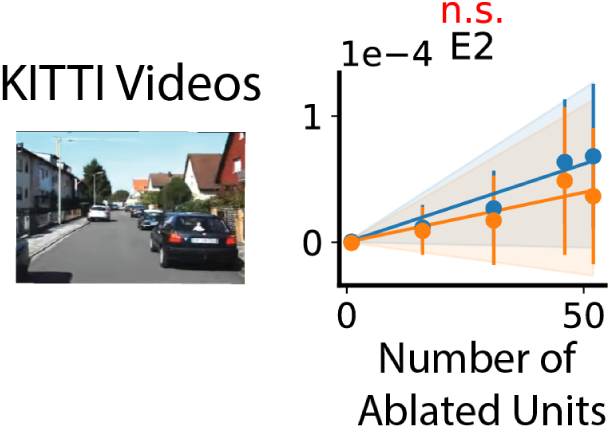
The effect of ablating the subsampled *E*_2_ BOS/non-BOS units on KITTI video prediction. Similar as Figure 4, for KITTI videos.

